# Microfluidic Extraction of Extracellular Vesicle-Associated Nucleic Acids for Downstream Molecular Analysis

**DOI:** 10.64898/2026.10.12.758106

**Authors:** Teresa Lage, Daniela Macedo, Hugo Silva, Alexandra Teixeira, Diogo Moreira, Natália Fiorenza, Ramón Lago Lestón, Ana Vilar Lagares, Belém Sampaio-Marques, Lara Pierantoni, Alejandro Rodriguez, Olga Azevedo, Paula Ludovico, Laura Muinelo Romay, Miguel Abal, Lorena Diéguez, Carlos Honrado, Alar Ainla

## Abstract

**Aim:** To develop a microfluidic workflow for extracting extracellular vesicle (EV)-associated nucleic acids (NAs) from biofluids, such as blood plasma and serum, for downstream molecular analysis.

**Methods:** Healthy serum and plasma samples were spiked with EVs obtained from Hec-1-A cells carrying defined mutations and characterized in accordance with MISEV guidelines. The spiked samples were then processed using either the standard centrifugation-based ExoGAG protocol or a microfluidic workflow. The latter combines ExoGAG mediated aggregation of EVs with mechanical entrapment of the resulting EV aggregates in a microfluidic filtration device (EVNA-chip). The retained material was washed and on-chip lysed to release its NA cargo. NAs were subsequently purified using a commercial silica column, producing an output suitable for molecular analysis through droplet digital PCR (ddPCR).

**Results:** The EVNA-chip recovery yield was 42.0 ± 4.28% and 38.1 ± 1.46% for double-stranded DNA (dsDNA) and microRNA respectively. The recovered dsDNA remained compatible with ddPCR detection of the KRAS p.G12D variant (c.35G>A) and with the assessment of MSI-associated alterations using BAT25, BAT26, NR21, NR24, and MONO27. The method was demonstrated for pilot cohort of patients with endometrial cancer. The RNA workflow supported reverse transcription and was demonstrated for the quantification of miR-X, a potential marker of Fabry disease, using patient samples and healthy controls.

**Conclusion:** Although recovery was lower than with the standard protocol, the centrifugation-free EVNA- chip workflow demonstrates the compatibility of the method with downstream NA molecular analyses and is advantageous for the development of an integrated, automated platform for EV-based liquid biopsy applications.

## INTRODUCTION

Liquid biopsy provides minimally invasive access to disease associated biomarkers and can support molecular detection, longitudinal monitoring, and treatment stratification^[1]^. Extracellular vesicles (EVs) are heterogeneous membrane bound particles continuously released by cells into extracellular spaces and biofluids^[2,3]^. Their molecular composition can reflect the physiological and pathological state of their cells of origin, providing a snapshot of the evolving genomic and transcriptomic landscape of diseases^[4]^. Importantly, a fraction of extracellular DNA and RNA is enclosed within or associated with extracellular vesicles^[5]^. Intraluminal nucleic acids are protected from extracellular nucleases by the vesicle membrane, which may enhance their stability relative to unprotected extracellular (cell free) nucleic acids^[6]^. These properties make EVs an attractive source of biomarkers for real-time molecular characterization, disease diagnosis, and monitoring^[7]^.

However, despite their clinical promise, translation of EV biomarkers remains limited by pre-analytical variability and by the absence of efficient, scalable, and clinically compatible EV isolation methods^[8,9]^. Differential ultracentrifugation is widely used but is time consuming, instrument intensive, and difficult to automate^[10]^. Size exclusion chromatography and filtration can improve standardization but dilute the sample and require additional pre-concentration^[11]^. Immunoaffinity methods offer molecular selectivity, yet they depend on chosen surface markers and may under-sample heterogeneous EV populations^[12]^.

Microfluidic technologies offer an attractive alternative by enabling sample processing in compact, low-volume devices that can be readily integrated into automated workflows ^[13]^. Most current microfluidic EV isolation platforms rely on immunoaffinity capture using antibodies directed against surface markers such as CD63^[14–19]^. While antibody-based approaches provide high specificity, they are associated with several limitations, including high cost, limited shelf life, and demanding storage requirements^[20]^. In addition, EV populations are heterogeneous, and not all vesicles express the same surface markers, potentially leading to biased recovery^[21]^. Chemical aggregation approaches, such as PEG-based (ExoQuick×) can overcome some of these limitations but can introduce polymeric contamination^[22]^. More recently, a precipitation method based on the interaction between EV surface sulfated sugars and the small molecule reagent dimethylmethylene blue (DMMB) has been developed and commercialized under the brand name ExoGAG^[23]^. ExoGAG exploits the interaction between a cationic agent and negatively charged, sulfated glycosaminoglycans (GAGs) present on the EV surface, inducing vesicle aggregation and facilitating their isolation^[23]^. The resulting aggregates can be recovered by low-speed centrifugation and have supported downstream analysis of DNA and RNA in biofluids^[24]^. Several studies have compared ExoGAG-mediated EV isolation with commonly used approaches, including 16 h ultracentrifugation (UC), size-exclusion chromatography (SEC), and commercially available precipitation kits such as ExoQuick and ExoSpin, across different biofluids, as conditioned medium, serum, plasma, urine and human milk. In Hec-1-A conditioned medium, Herrero et al. reported comparable particle recovery and significantly lower protein co-precipitation with ExoGAG than with 16 h ultracentrifugation^[25]^. In plasma, ExoGAG also reduced co-precipitated contaminants relative to the ExoQuick and ExoSpin precipitation methods without significantly compromising particle recovery^[24]^. In human milk, ExoGAG and ultracentrifugation provided the strongest performance for proteomic and transcriptomic analyses; however, caseins remained abundant in the isolated fractions, and the authors acknowledged that additional purification would be required for applications in which purity is critical ^[26]^. Overall, ExoGAG provides a simple, cost effective, and clinically compatible alternative to antibody-based enrichment strategies.

Here, we translate ExoGAG isolation protocol from a batch centrifugation process into a modular microfluidic method compatible with automation, providing a simple, low-cost platform for capturing EV aggregates and extracting their associated nucleic acids (Figure 1). Although several microfluidic approaches have addressed individual steps of EV isolation, release, lysis, or molecular detection, few integrate EV capture, washing, and lysis with subsequent nucleic acid recovery within a single workflow. Here the pillar-array mechanically retains EV aggregates, permits on-chip washing, and supports direct on-chip lysis, enabling efficient recovery of EV associated nucleic acids. Since the microfluidic device does not rely on surface functionalization it simplifies fabrication and subsequent storage and handling. We evaluate matrix dependent aggregation, nucleic acid recovery, and compatibility with downstream analysis using droplet digital PCR (ddPCR). The utility of the workflow for DNA analysis was examined using plasma and serum samples spiked with EVs derived from Hec-1-A cells, followed by ddPCR detection of KRAS p.G12D and a panel of microsatellite instability markers. These findings indicate that molecular alterations present in Hec-1-A cells can be detected in their released EVs and EV-spiked plasma or serum processed by our method. In a preliminary clinical demonstration, EV-associated DNA obtained from a patient with endometrial cancer yielded quantifiable microsatellite ddPCR signals. This method also supports RNA extraction, which we demonstrated by miR-X quantification in plasma samples from individuals with Fabry disease and healthy controls. Lower miR-X levels were observed in the Fabry disease group. Collectively it shows that this simple and tunable EV-associated NA isolation method, which would be compatible with future automation, supports downstream analysis of NA recovered from complex biofluids. This study therefore provides a technical foundation for further development of EV associated NA analysis for minimally invasive liquid biopsies.

**Figure 1.**
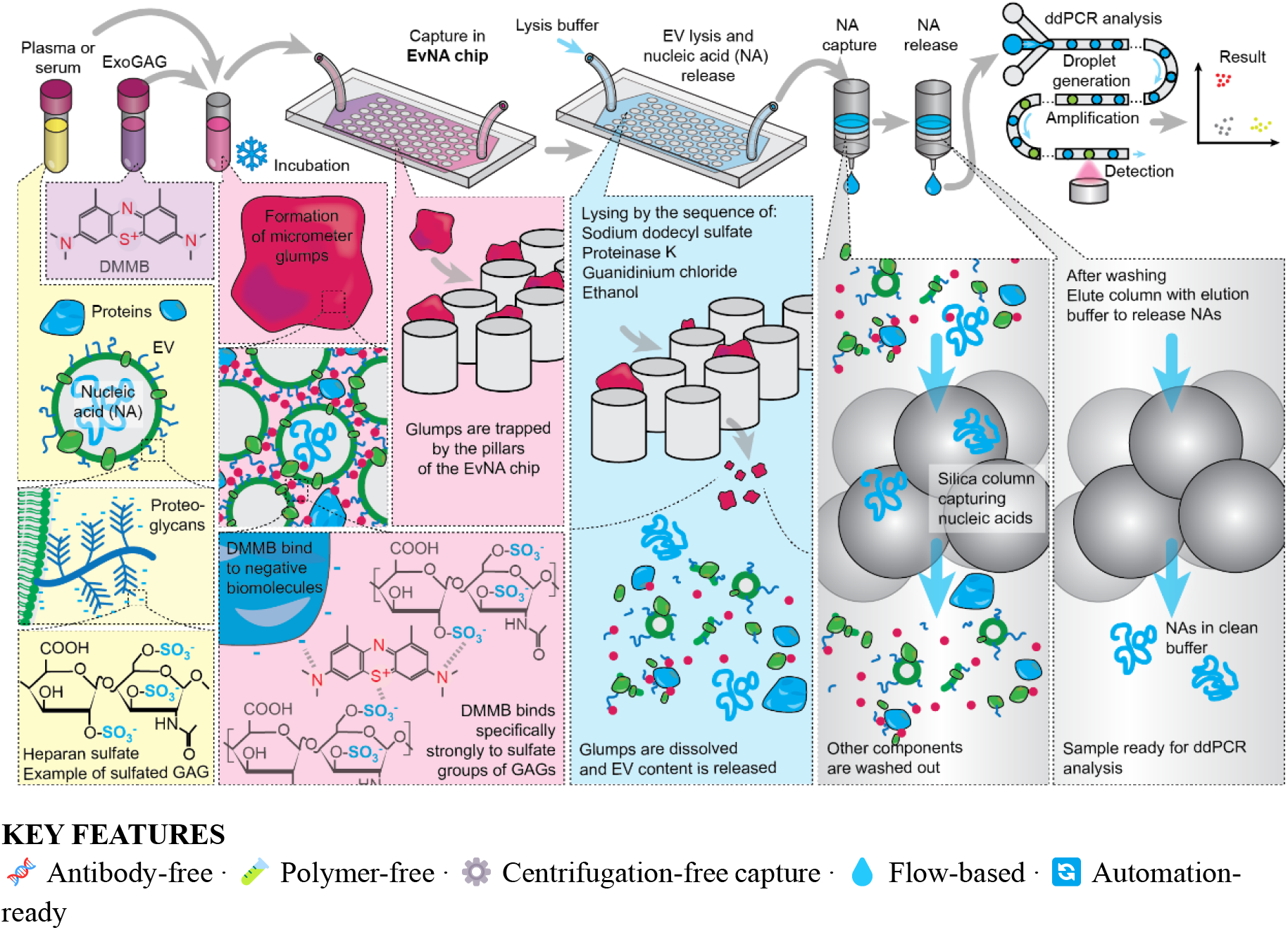
Microfluidic method for the EV nucleic acids analysis from plasma or serum. Method steps (above) and corresponding conceptual mechanisms (below). DMMB binding to sulfated GAGs on EV surface and other biomolecules form microscopic precipitates, which are mechanically trapped by the pillars in the EVNA-chip. Lysing releases the EV cargo, which is processed through silica column capturing nucleic acids. After elusion pure nucleic acid sample is obtained and quantified using ddPCR.

## METHODS

### Materials

ExoGAG (Nasasbiotech, A Coruña, Spain) was used to induce the aggregation of EV-containing material, Microfluidic devices were fabricated in polydimethylsiloxane (PDMS) and operated using syringes, tubing, connectors and a syringe pump. DNA was purified using a commercial silica column and the QIAamp DNA mini kit (QIAGEN, Venlo, the Netherlands), whereas total RNA was isolated using the Plasma/Serum Exosome RNA Isolation Kit (product 58000; Norgen Biotek). DNA and total RNA were quantified using the Qubit dsDNA High Sensitivity and microRNA High Sensitivity Assay Kits, respectively (Thermo Fisher Scientific). Reverse transcription and ddPCR analysis were performed using the TaqMan MicroRNA Reverse Transcription Kit, a miR-X specific TaqMan MicroRNA Assay, ddPCR Supermix for Probes, KRAS p.G12D mutant- and wild type specific assays, and the Bio-Rad MSI multiplex assay.

### Device design and fabrication

Microfluidic devices were fabricated in polydimethysiloxane (PDMS) using a soft lithographic technique. First the circuitry was designed in AutoCAD (Autodesk, San Francisco, CA, USA). The EVNA-chip comprises six parallel chambers, each containing two sequential pillar-array regions. The first region contains 120 µm diameter pillars separated by 20 µm gaps, while the second contains 130 µm-diameter pillars separated by 10 µm gaps (Figure 3). The chip has an internal volume of approximately 33.9 µL and contains approximately 90,000 pillars.

For plasma samples, two upstream pre-chips with pillar gaps of 600 and 270 µm, respectively, and a structure height of 250 µm for both chips were used sequentially (EVNA-prechip). The modular design also allows multiple chips to be connected in series and configured for different sample types. Master molds were fabricated in INL class 100 cleanroom facility. The designed pattern was transferred to a photoresist on a silicon wafer using direct laser writing (DWL 200, Heidelberg Instruments). After development, silicon was etched by deep reactive ion etching (DRIE, SPTS Pegasus) to obtain desired channel height. After etching, C_4_F_8_ was plasma polymerized to form non-stick coating. Then the master molds were used to cast Sylgard 184 (Dow Corning, Midland, MI) 10:1 PDMS pre-polymer mixture and cured in the oven (at 65°C for 3h). Cured PDMS structure was released, fluid ports were punched, and the device was plasma bonded to a glass microscopy slide using oxygen plasma cleaner (Harrick Plasma, Ithaca, NY). Completed devices were stored in ambient conditions until use

### EVs sample preparation

#### Cell culture and conditioned medium preparation

The human endometrial cancer cell line Hec-1-A was cultured in McCoy’s 5A media (Gibco, Grand Island, NY, USA) supplemented with 1% penicillin and streptomycin (Gilco, Grand Island, NY, USA) and 10% FBS (Gilco, Thermo Fisher Scientific) and maintained in a humidified incubator with 5% CO2 at 37ºC. The cell line was routinely tested and confirmed mycoplasma contamination-free. EVs collection was performed after culturing cells for 48h with medium with 10% of FBS reduced in EVs.

#### Ultracentrifugation (UC)

Conditioned medium (20 mL) was sequentially centrifuged at 500 × g for 10 min and 2,000 × g for 10 min to remove cells and cellular debris. The resulting supernatant was ultracentrifuged at 100,000 × g for 16 h at 4 °C using a 70 Ti rotor (Beckman Coulter, Brea, CA, USA). The EV-enriched pellet obtained after this step was resuspended in Phosphate-buffered saline (dPBS) to a final volume of 20 mL and subjected to a second ultracentrifugation at 100,000 × g for 2 h at 4 °C. The final EV-enriched pellet was resuspended in dPBS and stored at ™80 °C until further use.

### Characterizations of EVs

#### Transmission electron microscopy (TEM)

EVs were isolated from 20 mL of conditioned medium (CM) collected from Hec-1-A cell cultures. The CM was processed according to the isolation protocol described above and subsequently concentrated by ultrafiltration to a final volume of 500 µL. The concentrated EV preparation was then further purified by size-exclusion chromatography (SEC) using qEVoriginal 70 nm Gen 2 columns (Izon, Christchurch, New Zealand), following the manufacturer’s instructions. dPBS was used as the eluent, and four fractions of 400 µL each were collected. To further enrich the EV preparation, the pooled SEC fractions were concentrated by ultrafiltration to a final volume of 50 µL. Before grid preparation, the sample was diluted 1:1 with Milli-Q® water.

For transmission electron microscopy (TEM) analysis, Quantifoil holey carbon grids (Quantifoil, Jena, Germany) were plasma-treated under high vacuum for 4 s using a Fischione 1020 Plasma Cleaner (Fischione Instruments, Pennsylvania, USA) to increase grid hydrophilicity and minimize surface contamination. A 5 µL aliquot of the EV suspension was deposited onto each grid and allowed to adsorb for 2 min. Excess liquid was carefully removed using filter paper. Negative staining was performed by applying 2% uranyl acetate twice, with each staining step lasting 30 s.

The stained grids were examined using a JEOL JEM-2100 transmission electron microscope (JEOL Ltd., Akishima, Tokyo, Japan) equipped with a LaB_6_ filament and operated at an accelerating voltage of 200 kV. Images were acquired at magnifications ranging from 40,000× to 80,000×, enabling detailed morphological and structural characterization of the isolated EVs.

#### Nanoparticle Tracking Analysis (NTA)

Size distribution and concentration of EVs was assessed by nanoparticle tracking analysis (NTA) using Malvern Panalytical NanoSight NS300 (Malvern Instruments, Malvern, UK), configured with a Blue 488nm laser and a high sensitivity sCMOS camera. The absence of background was verified using 0.2 µm filtered Milli-Q water. For NTA analysis dPBS was used as a diluent, and five videos of 60 s were captured at camera level 14. The videos were then analyzed using the NTA 3.0 software version (Malvern Instruments, Malvern, UK) with a detection threshold of 10.

#### Western Blot Analysis (WB)

Total EV protein was extracted using 2.5% w/v urea prepared in Milli-Q water and supplemented with Phenylmethylsulfonyl fluoride (PMSF), Protease Inhibitor Cocktail (PIC) and protein extracts were quantified by bicinchoninic acid assay (Pierce™ BCA Protein Assay, thermofisher). 20µL protein lysates were mixed with 4× Laemmli sample buffer without reducing agent and heated at approximately 100 °C for 10 min prior to electrophoretic separation on SDS-PAGE gels. SeeBlue™ Plus2 Pre-Stained Protein Standard (Thermo Fisher Scientific) was used as a molecular weight marker. Following electrophoresis, proteins were transferred onto PVDF membranes by wet transfer at 100 V for 90 min. Membranes were subsequently blocked with 5% non-fat milk prepared in PBS-T for 1 h at room temperature and then incubated overnight at 4 °C with primary antibodies against CD63 (1:500, Santa Cruz), flotillin (1:500, Proteintech), Alix (1:500, Thermo Fisher Scientific) and calnexin (1:5000, Abcam).

After washing, membranes were incubated for 1 h at room temperature with the appropriate HRP-conjugated secondary antibodies (goat anti-mouse or goat anti-rabbit) diluted 1:5000 in PBS-T. Immunoreactive bands were detected using SuperSignal™ West Femto Chemiluminescent Substrate, prepared at a 1:1 ratio according to the manufacturer’s recommendations.

Chemiluminescent signals were visualized using a Syngene G CHEMI XT4 imaging system. For each membrane, two images were acquired with exposure times of 30 s to ensure optimal signal detection.

### On-chip EVs isolation and nucleic acids recovery

For the plasma and serum experiments, 50 µL of HEC-1-A-derived EVs, isolated using the 16 h + 2 h ultracentrifugation protocol described in detail above, was spiked into 500 µL of serum or plasma and incubated with 1,100 µL of ExoGAG (Nasasbiotech, A Coruña, Spain) for 5 min. Resulting mixture was passed through series of EVNA chips at flow rate 20 µL min^−1^. In case of serum 2× EVNA-chips were used. Plasma produced larger and more abundant aggregates than serum; therefore, the plasma workflow incorporated further two upstream EVNA-prechips followed by two standard capture modules (2× EVNA- chip). Captured material was lysed by introducing 180 µL sodium dodecyl sulfate solution containing 20 µL proteinase K at 70 °C. Flow was applied for 3 min at 50 µL min^−1^ followed by 7 min at 20 µL min^−1^. This was followed by 200 µL guanidinium chloride solution at 70 °C using the sequence of same flow rates. The combined lysate was collected, mixed with 200 µL ethanol at room temperature, transferred to a silica column (QIAGEN, Hilden, Germany), washed according to the procedure of the commercial kit, and eluted in 100 µL of buffer Tris-EDTA (TE). DNA was quantified using the Qubit high-sensitivity dsDNA assay.

The capture protocol was unchanged for RNA experiments. Captured material was lysed at room temperature using 300 µL guanidinium thiocyanate solution supplemented with 37.5 µL mild non-ionic detergent. The solution was introduced for 3 min at 50 µL min^−1^ and 7 min at 20 µL min^−1^. The lysate was collected, mixed with 500 µL ethanol, and processed using the Exosomal RNA Isolation Kit (Norgen Biotek). RNA was eluted in 25 µL of Buffer TE and quantified using the Qubit RNA high-sensitivity assay.

### Standard ExoGAG isolation protocol

Serum or plasma (500 µL) was incubated with 1000 µL of ExoGAG (Nasasbiotech, A Coruña, Spain) EVs precipitation formulation for 5 min at 4ºC. After incubation, the mix was centrifuge at 3000 × g, 20 min, at 4 ºC. The supernatant fraction was discarded, and the pellet was resuspended in dPBS.

### DNA and small RNA extraction and measurement

DNA was extracted from plasma samples using the QIAamp DNA Mini Kit (QIAGEN, Hilden, Germany), following the manufacturer’s instructions.

Total RNA was extracted from previously isolated EVs using the Plasma/Serum Exosome RNA Isolation Kit (Norgen) (Product 58000), following the manufacturer’s instructions.

All extractions were performed using the same volume of plasma or serum (500 µL) and the concentration was normalized based on the extraction volume and the elution volume. The quantification of DNA and microRNA were performed using NanoDrop Spectrophotometer and Qubit 4.0 Fluorometer with the Qubit™ dsDNA High Sensitivity (HS) Assay Kit and Qubit™ microRNA Assay Kit, respectively, according to the manufacturer’s instructions

### Reverse Transcription of miR-X

A TaqMan MicroRNA Assay (Applied Biosystems) was used to assess miR-X abundance in samples from patients with Fabry disease. Briefly, 10 ng of total RNA recovered from EV containing preparations was reverse transcribed using miR-X specific stem-loop reverse transcription primers and the TaqMan MicroRNA Reverse Transcription Kit (Applied Biosystems), according to the manufacturer’s instructions. Reverse transcription was performed using a T100 Thermal Cycler (Bio-Rad Laboratories).

### Droplet digital PCR (ddPCR)

DNA was analyzed using the QX200 Droplet Digital PCR System (Bio-Rad). The KRAS p.G12D variant (c.35G>A) was detected using fluorescent probes specific to mutant and wild-type alleles. MSI associated alterations were analyzed in the extracted DNA using the Bio-Rad MSI multiplex assay on the QX200 Droplet Digital PCR System (Bio-Rad, Hercules, CA, USA) with TaqMan probe chemistry. Five microsatellite markers were assessed in three multiplex reactions: BAT25/BAT26, NR21/NR24, and MONO27. The assay uses two competing probes targeting the same microsatellite sequence. Changes in repeat length alter the binding of the probes, allowing shortened and wild-type alleles to be distinguished when their length differs by at least two base pairs. Samples were analyzed in duplicate. Droplets were generated using the QX200 Automated Droplet Generator according to the manufacturer’s instructions. PCR amplification was performed in a C1000 Touch Thermal Cycler (Bio-Rad) under the following conditions: 95°C for 10 min, followed by 40 cycles of 95°C for 15 s and 55°C for 60 s, and a final step at 98°C for 10 min. A ramp rate of 2.5°C/s was used throughout the protocol. Droplets were read using the QX600 Droplet Reader, and the data were analyzed with QuantaSoft software. Droplets were classified as mutant, wild-type, or empty according to their fluorescence signal. A sample is considered microsatellite instability high (MSI-H) when at least two of the five markers are positive^[27]^.

RNA recovered from EV-spiked plasma processed by our microfluidic workflow was reverse transcribed using a target-specific procedure for miR-X. The resulting cDNA was subsequently analyzed using a ddPCR protocol adapted for miR-X detection. Each 22 µL reaction contained 10 µL of 2× ddPCR Supermix for Probes (No dUTP; Bio-Rad Laboratories), 1 µL of the miR-X specific TaqMan MicroRNA Assay (Applied Biosystems), 6 µL of nuclease-free water and 5 µL of cDNA. For the no-template control (NTC), the cDNA was replaced with an equal volume of nuclease-free water. Reaction mixtures were transferred to a 96-well plate, which was sealed using foil at 180°C for 5 s. The plate was then vortexed vigorously and briefly centrifuged at 1150 × g. Droplets were generated using the Automated Droplet Generator (Bio-Rad Laboratories) according to the manufacturer’s instructions. PCR amplification was performed using the following thermal cycling conditions: enzyme activation at 95°C for 10 min, 45 cycles of denaturation at 94°C for 30 seg and annealing at 55°C for 1 min, enzyme deactivation at 98°C for 10 min, and a final hold at 4°C for 30 min. Fluorescence data were acquired using the QX600 Droplet Reader (Bio-Rad Laboratories). To assess the compatibility of the recovered RNA with downstream miRNA analysis, plasma samples from healthy controls and patients with Fabry disease were analyzed alongside the corresponding positive and negative controls. All samples and controls were analyzed in duplicate.

### Study participants and sample collection

Endometrial cancer cohort: A total of eleven blood samples were collected: seven patients with endometrial cancer (EC) were recruited between January 2018 and June 2021 at the Gynecology Department of Vall d’Hebron University Hospital (Barcelona, Spain), MD Anderson Cancer Center (Madrid, Spain), and University Clinical Hospital of Santiago de Compostela (Santiago de Compostela, Spain) and four from healthy individuals. Patients were eligible for inclusion if they met the following criteria: (i) diagnosis of endometrial carcinoma of any histological subtype; (ii) age ≥18 years; (iii) provision of written informed consent; (iv) known mismatch repair (MMR) protein status; (v) absence of ongoing antitumor treatment at the time of sample collection; and (vi) no history of another malignant tumor during the five years preceding sample collection. Peripheral blood was collected into CellSave Preservative Tubes (Silicon Biosystems Inc., Huntingdon Valley, PA, USA). Plasma was isolated using a two-step centrifugation protocol.

Fabry disease cohort: A total of six peripheral blood samples were collected: three from individuals with a confirmed diagnosis of Fabry disease (FD) and three from healthy individuals. The FD cohort included: (i) asymptomatic individuals; (ii) patients with mild-to-moderate cardiac involvement, defined as left ventricular hypertrophy without late gadolinium enhancement or dysrhythmias; and (iii) patients with renal involvement, defined by albuminuria category A2 or A3. Exclusion criteria were: (i) age <18 years; (ii) neurological involvement attributable to FD; (iii) detectable antibodies against agalsidase alfa or agalsidase beta; and (iv) ongoing immunosuppressive treatment. Samples were collected at the Reference Center for Lysosomal Storage Disorders, Hospital Senhora da Oliveira, Guimarães (ethics approval reference 74/2021), and at the Clinical Academic Center–Braga (2CA-Braga; ethics approval reference 37_2022), Portugal. Samples were processed at the Life and Health Sciences Research Institute, University of Minho. All participants provided written informed consent. The study protocol was approved by the National Provisional Committee on Ethics in Health Research and conducted in accordance with the principles of the Declaration of Helsinki. Peripheral blood (10 mL) was collected by venipuncture using heparin as an anticoagulant and immediately processed. Plasma was isolated using a two-step centrifugation protocol.

### Statistical analysis

Nucleic acid recovery was calculated as 100 × (amount recovered after on-chip lysis / amount introduced). Final dsDNA and microRNA recovery values are reported as mean ± the dispersion reported in the source dataset for three replicate experiments.

## RESULTS

In the presented work, our aim has been to develop a simple and practical method for the isolation of NA cargo of extracellular vesicles (EVs) from biofluids. This method shall be implementable into compact (e.g. desktop sized) automated and cost-effective instrument, overcoming current limitations of large centrifuges and manual handling. Besides instrumentation, other important practical consideration is the handling of reagents and consumables, which should be stable and have long shelf-life. Therefore, we have chosen to build upon chemical precipitation of EVs using ExoGAG, which is commercially available formulation specifically designed for EVs isolation. The main active ingredient of ExoGAG is 1,9- dimethyl-methylene blue (DMMB) ^[23]^. DMMB is a well-established metachromatic dye used for histological staining and the quantification of sulfated glycosaminoglycans (sGAGs) ^[28,29]^, binding to various negatively charged biomolecules and has specifically high affinity to sulfated glycosaminoglycans (GAGs) on the membranes of cells and EVs. At the assay pH of 3.5, which is below the isoelectric point of many proteins, these proteins tend to carry a net positive charge, reducing their electrostatic interaction with the cationic DMMB, whereas sulfate groups remain negatively charged. However, other polyanionic biomolecules, such as DNA, RNA, and hyaluronic acid, may also interact with the DMMB under acidic conditions, limiting its specificity. Compared to antibodies used for EVs capture, DMMB is less specific, targeting all EVs and it has high stability in solution. Previous work describing the ExoGAG method report that DMMB co-precipitate approximately 15% of the cell free DNA (cfDNA) present in the sample ^[23]^. These observations were interpreted as suggesting that most of the recovered DNA using ExoGAG was associated with EVs rather than derived from free cfDNA^[23]^. This limited cfDNA remnant, together with the successful detection of DNA targets after processing, is consistent with the hypothesis that at least part of the recovered DNA is associated with vesicles or other structures that preserve nucleic-acid integrity. The standard ExoGAG procedure for EVs isolation still requires conventional low-speed centrifuge to collect the aggregates. While accelerating and simplifying the manual workflow compared to the ultracentrifugation, it is still not easy to fully automate. Here we have translated this standard procedure into microfluidic format, using flow through chips and columns without any need for centrifugation. EV aggregates would be captured in EVNA-chips, which are composed of arrays of precisely spaced pillars. Depending on the sample several EVNA-chips could be connected even in series. By changing the input solution, captured aggregates can be washed, and EVs lysed directly in the EVNA-chip, after which released NAs can be isolated using commercially available silica column, from which the clean eluted sample would be ready for NA detection (e.g. using PCR). We have optimized the chip design and operation protocol and characterized the performance of EVNA-chips to isolate and retrieve NA from EVs in different common sample matrixes, such as plasma and serum and demonstrate that resulting material is suitable for sensitive detection using ddPCR.

### EV model samples

To enable reproducible testing of microfluidic devices, first we established a standardized EV sample, which were based on spiking (50 µL) of concentrated EVs into sample matrices (500 µL), such as serum or plasma (Figure 2). A standardized Hec-1-A EV-enriched preparation purified from conditioned medium collected from Hec-1-A cell cultures using standard UC based method and characterized following MISEV 2023 Guidelines for EVs studies using NTA, TEM and Western blotting to detect EV-associated proteins ALIX, CD63 and Flotillin-1, confirming presence of EVs in the preparation. Furthermore, because the molecular profile of the Hec-1-A cell line is well characterized, its known molecular alterations were used as traceable targets to determine whether the genetic material recovered using the microfluidic device remained suitable for downstream molecular analyses ^[30]^.

**Figure 2.**
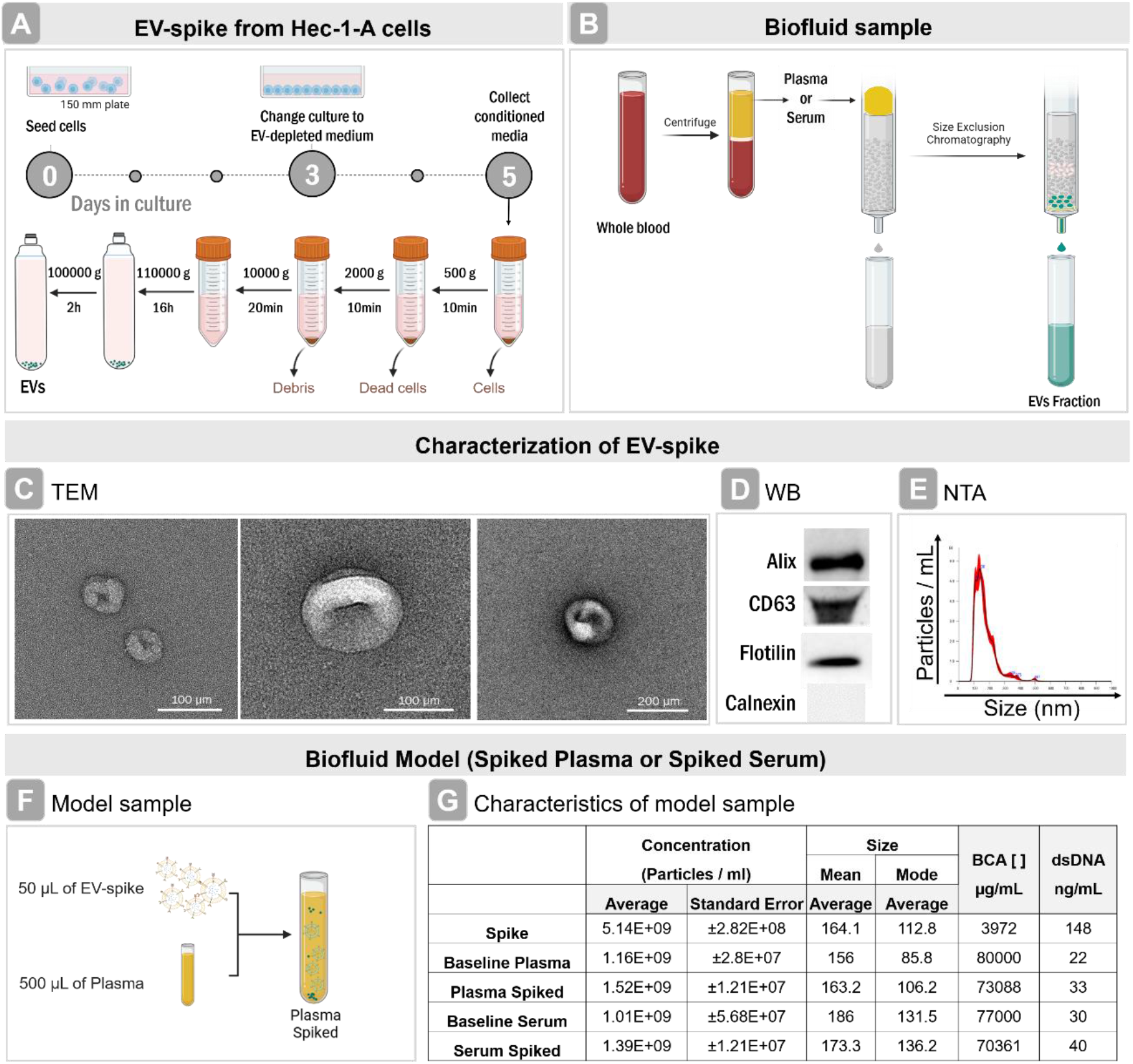
Preparation and characterization of a standardized EV model sample. (A) Preparation of EV- spike from Hec-1-A cell conditioned medium (B) Isolation of EVs from biological matrix. Characterization of EV-spike by (C) transmission electronic microscopy (TEM), (D) Western-blotting for the EV-associated proteins ALIX, CD63, flotillin-1, and the cellular contamination marker calnexin (E) nanoparticle tracking analysis (NTA). (F) Formulating models samples by introducing EV-spike into sample matrix (plasma or serum), (G) Characteristics of model samples and their components for particle concentration, size, protein and nucleic acid content.

**Figure 3.**
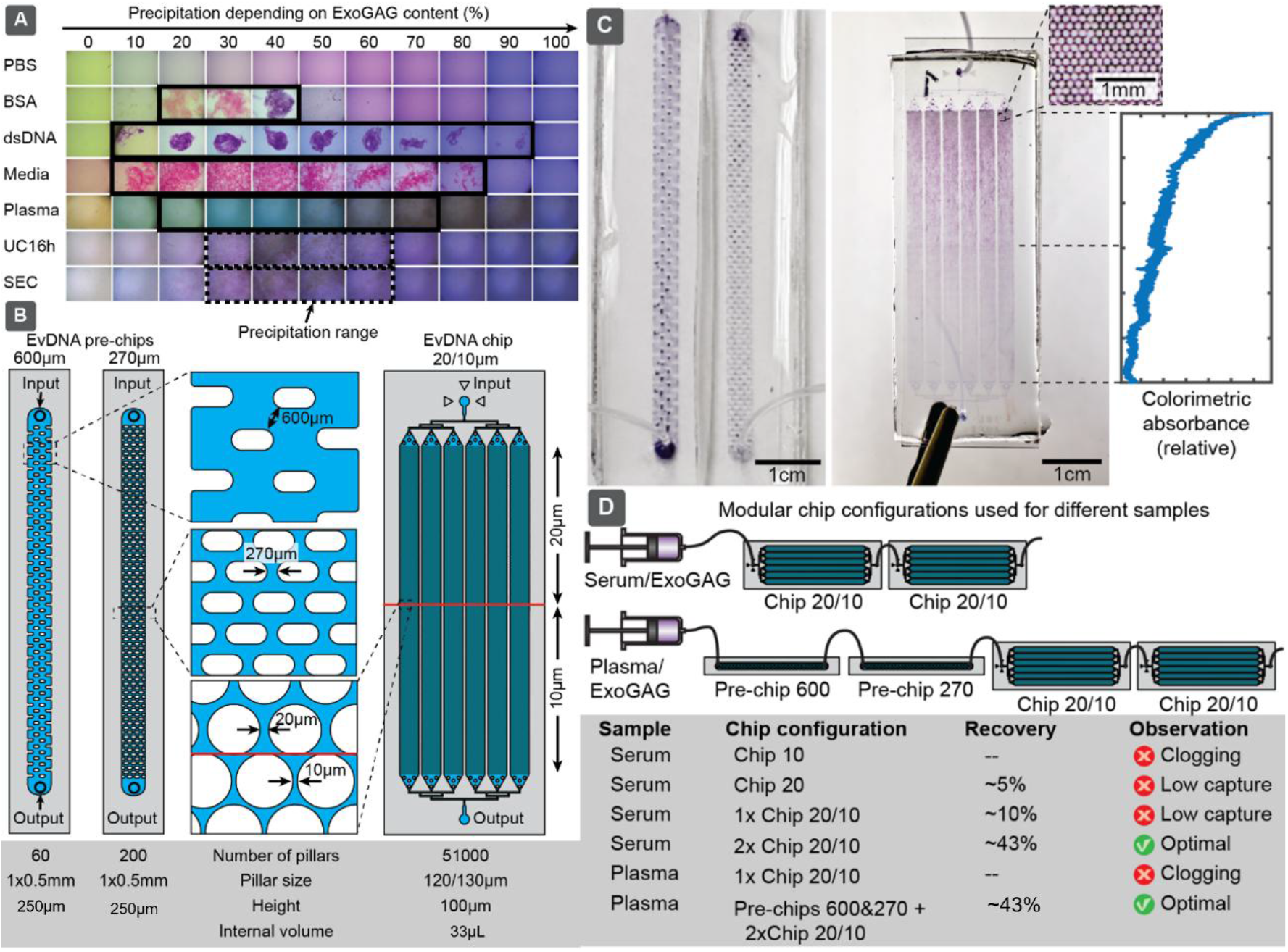
Matrix-dependent ExoGAG aggregation and EVNA pillar-chips to capture them. (A) Visible aggregate formation following the mixing of ExoGAG with different biological matrices. (B) Schematic representation of three EvNA chip designs featuring gaps of 600, 270 and 20/10 µm. (C) Photographs of EVNA chips loaded with ExoGAG induced aggregates. Colorimetric graph shows relative distribution of the aggregates along the pillar array. (D) Modular configurations of chips connected in series used to process serum and plasma samples.

These model samples were used throughout the study. Purpose of spiking was not to significantly alter the overall matrix composition, but to introduce Hec-1-A specific nucleic acid sequences, that can be detected at the output. Also, final formulations were analyzed using size-exclusion chromatography (SEC) and obtained concentrations followed expected values (Figure 2G).

Compared to the corresponding baseline biofluids, the spiked samples retained a similar overall particle size distribution while showing a modest increase in particle concentration. Total protein concentration decreased slightly following spiking, whereas dsDNA concentrations increased (Figure 2G). This limited change was consistent with the relatively low protein and dsDNA content of the spike-in preparation compared with the baseline levels present in serum or plasma. The model was designed to produce a moderate increase in particle abundance, approximating the elevated circulating particle levels reported in certain pathological conditions, including some cancers, without substantially altering the composition of the original biofluid.

### Influence of the matrix composition to ExoGAG induced aggregation

As the underlying principle of EVs capture in our method relies on mechanical trapping of aggregates, we first investigated qualitatively the precipitation process of different samples and biological components depending on the added ExoGAG content. In our observations, the aggregate formation depended on both, the sample composition and its relative proportion in the ExoGAG mixture (Figure 3A). In figure 3A, each row represented a different matrix, while the columns showed a compositional gradient from ExoGAG alone to sample alone, in 10% increments. As expected, PBS did not form visible precipitates with ExoGAG. In contrast, dsDNA (∼ 20 kb) was precipitated by any amount of ExoGAG. Media (McCoy 5A) and BSA solution also formed aggregates with ExoGAG, showing that ExoGAGs binding to both proteins and nucleic acids. Serum and Plasma formed visible aggregates predominantly at higher ExoGAG proportions, ranging from 50% to 90%, with plasma producing the largest amount of visible aggregates among the biofluids tested, due to the highest concentration of proteins. Purified EV samples obtained by either 16 h ultracentrifugation or size-exclusion chromatography also formed visible aggregates particularly, when ExoGAG content was 30%–60%. Collectively, these qualitative observations indicate that ExoGAG-mediated aggregation is strongly influenced by the biochemical composition of the sample and the relative proportions of ExoGAG added, but also that ExoGAG not only bind to EVs but at different degree to various biological macromolecules and nanoparticles, which agrees with previous studies, that have shown, that while DMMB has stronger affinity to sulfated-GAGs it also binds other biomolecules ^[28,29]^. Furthermore, we complemented the visual observation also with a centrifugation experiment, comparing standard ExoGAG protocol applied to plasma and serum with purified EV samples. In case of plasma 3,000× g for 20min was sufficient for the sedimentation, however this was not enough to pellet aggregates from purified EVs in dPBS, where centrifugation speed needed to be increased to 16,000× g to achieve similar recovery, also confirmed by complete recovery of DNA. In case of plasma sample 16,000× g protocol produced highly compact pellet that were difficult to resuspend. In case of similar viscosity and densities, sedimentation speed in centrifugation shall be proportional to the g-force and cross-section of spherical particles (*v* ∝ *gr*^2^), thus suggesting, that EVs alone form smaller aggregates compared to the biological matrix, where other biomolecules could also co-precipitate and facilitate formation of larger aggregates. These variations in precipitation guide also fluidic design, which needed to achieve high capture efficiency while avoiding clogging.

### Microfluidic EV capture and nucleic acid recovery

The EVNA-chips were designed as a mechanical filter for the retention of EV-containing ExoGAG aggregates. Different inter-pillar gaps were evaluated to identify the most suitable device configuration (Figure 3). Overall, the microfluidic design needed to address two main criteria: i) being able to hold target sample, and ii) being able to capture aggregates with high yield. Too small gaps would result in device clogging and mechanical failure before entire sample is processed, whereas too large would result in low capture efficiency. In desired configuration precipitates would be distributed homogeneously across the pillar array of the chip. To achieve the optimal capture, we found that microfluidic design shall contain regions with different gap sizes. These can be implemented in the same chip as shown on figure 3B, or microfluidic design can be also made modular, where chips with same or different gap sizes can be connected in series to address flexibly different samples. Here the eventually chosen configuration comprised three chip designs: i) EVNA-chip, which had two sequential pillar-array regions with different inter-pillar gap sizes. The first region contained pillars with a diameter of 120 µm and gaps of 20 µm, whereas the second contained 130 µm pillars separated by 10 µm gaps. This sequential configuration was selected to retain aggregates of different sizes, while balancing capture capacity, hydrodynamic resistance, fabrication feasibility and mechanical stability. Pillar geometries with smaller cross-sections and higher aspect ratios were excluded because they were prone to fracture during demolding; and ii) two EVNA- prechips with inter-pillar gaps of 600 and 270 µm, respectively, and a structure height of 250 µm for both chips were (Figure 3). These chips were connected before the main EVNA-chip in case of sample, which would produce very larger aggregates.

To process 500 µL of EV-spiked serum sample and maximize the retention we connected two EVNA-chips in series and processed sample at flow rate of 20 µL/min. Same amount of plasma sample, however, produced larger and more abundant aggregates than serum, which clogged the device configuration used for serum. Therefore, plasma samples were processed through series of four chips, where two EVNA- prechips (with 500 and 250µm gaps) were added before 2 × EVNA-chips.

Following retention, the captured material was lysed directly within the devices using the optimized SDS, proteinase K and guanidinium chloride protocol. The lysate was collected and transferred to a commercial silica column. Purified DNA was eluted in 100 µL of buffer TE and quantified using a Qubit fluorometer. Recovery efficiency was calculated by comparing the amount of nucleic acid initially introduced into the system with the amount recovered after on-chip capture, lysis and purification.

For the serum experiments, 50 µL of the concentrated Hec-1-A EV-enriched preparation was added to 500 µL of serum and mixed with 1.1 mL of ExoGAG. The concentrated preparation contained an average of 19.0 ± 4.6 ng of EV-associated DNA. The standard ExoGAG protocol recovered 93.6 ± 9.4% of the initial EV-associated dsDNA, whereas the microfluidic workflow recovered 42.0 ± 4.28% of the input double-stranded DNA (n = 3). In case of plasma microfluidic system recovered 40.7 ± 2.28% of the double-stranded DNA (n = 3).

In the RNA experiments, the concentrated EV-enriched preparation had an average of 28.5 ng of EV-associated microRNA. The standard centrifugation-based ExoGAG protocol recovered 93.2% of the input microRNA. By comparison, the microfluidic workflow recovered 38.1 ± 1.46% of the available microRNA (n = 3).

Although in general recovery was lower than that obtained with the standard protocol, the dsDNA and RNA collected after on-chip lysis remained suitable for downstream ddPCR analysis.

### Detection of KRAS G12D and MSI in DNA from spiked Plasma and Serum

KRAS p.G12D is a clinically relevant point mutation that may inform therapeutic stratification in selected cancer settings. In this study, it was used as a traceable molecular target because the mutant signal was detected in Hec-1-A cellular DNA. The KRAS p.G12D signal was subsequently detected in DNA recovered from Hec-1-A EV-enriched preparations. It was also detected in EV-spiked serum and plasma processed either by the microfluidic workflow or by the standard ExoGAG protocol (Figure 4). Across all tested conditions, the mutant and wild-type droplet clusters remained distinguishable following on-chip lysis and silica-column purification, demonstrating that the recovered DNA remained suitable for variant-specific ddPCR analysis.

**Figure 4.**
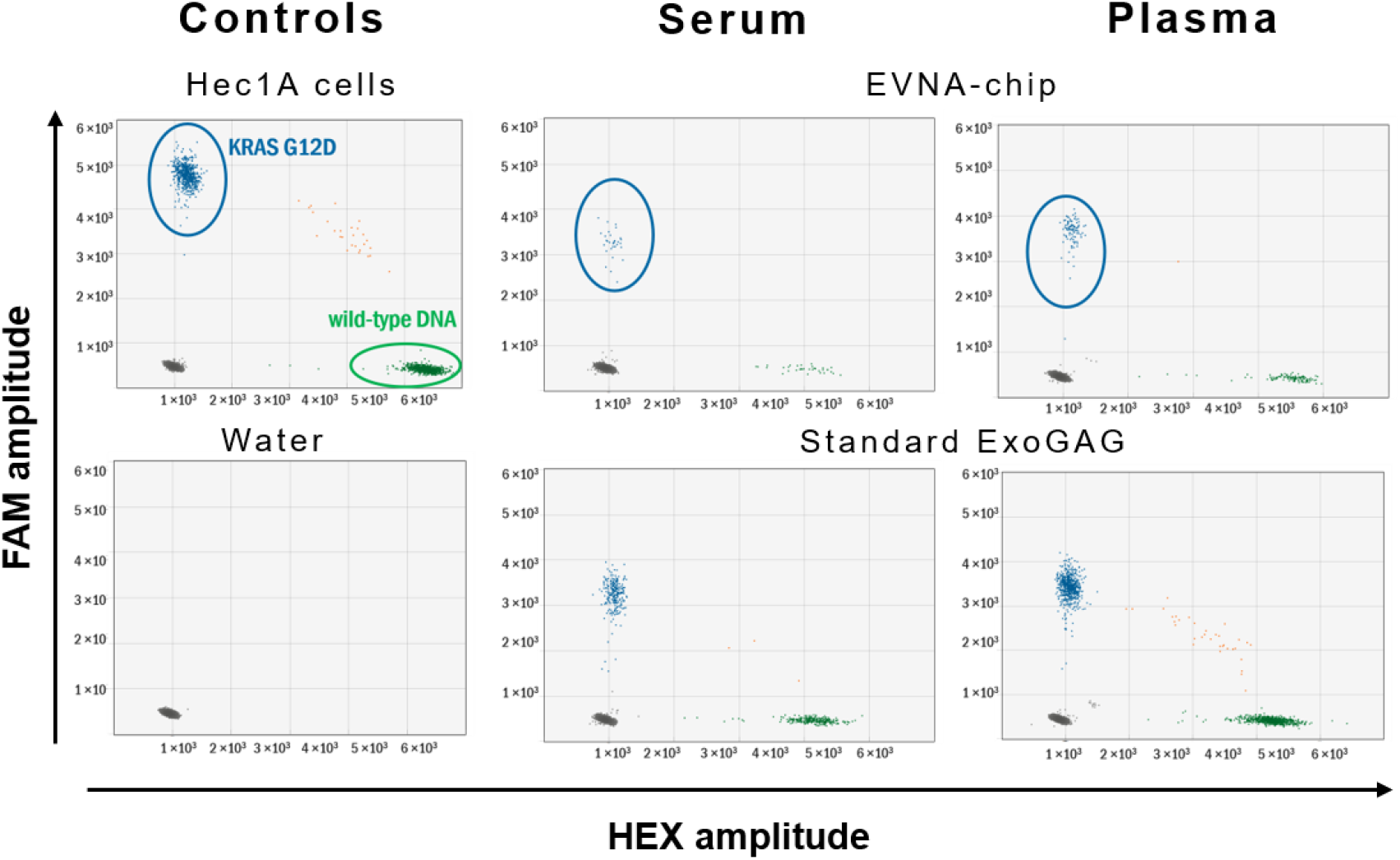
Detection of KRAS p.G12D by ddPCR in DNA recovered using the standard ExoGAG protocol and the EVNA-chip workflow.

MSI-associated alterations were evaluated using a multiplex ddPCR assay targeting five mononucleotide repeat markers across three reactions: BAT25/BAT26, NR21/NR24 and MONO27. The known microsatellite alterations of the Hec-1-A cell line were used as a positive analytical model. MSI-associated ddPCR signals were detected in DNA recovered from the Hec-1-A EV-enriched preparation and from Hec- 1-A EV-spiked serum and plasma processed using either the EVNA-chip workflow or the standard ExoGAG protocol (Figure 5). Marker-specific signals remained detectable after on-chip capture, lysis and DNA purification, showing that the recovered DNA was also suitable for downstream MSI-associated ddPCR analysis.

**Figure 5.**
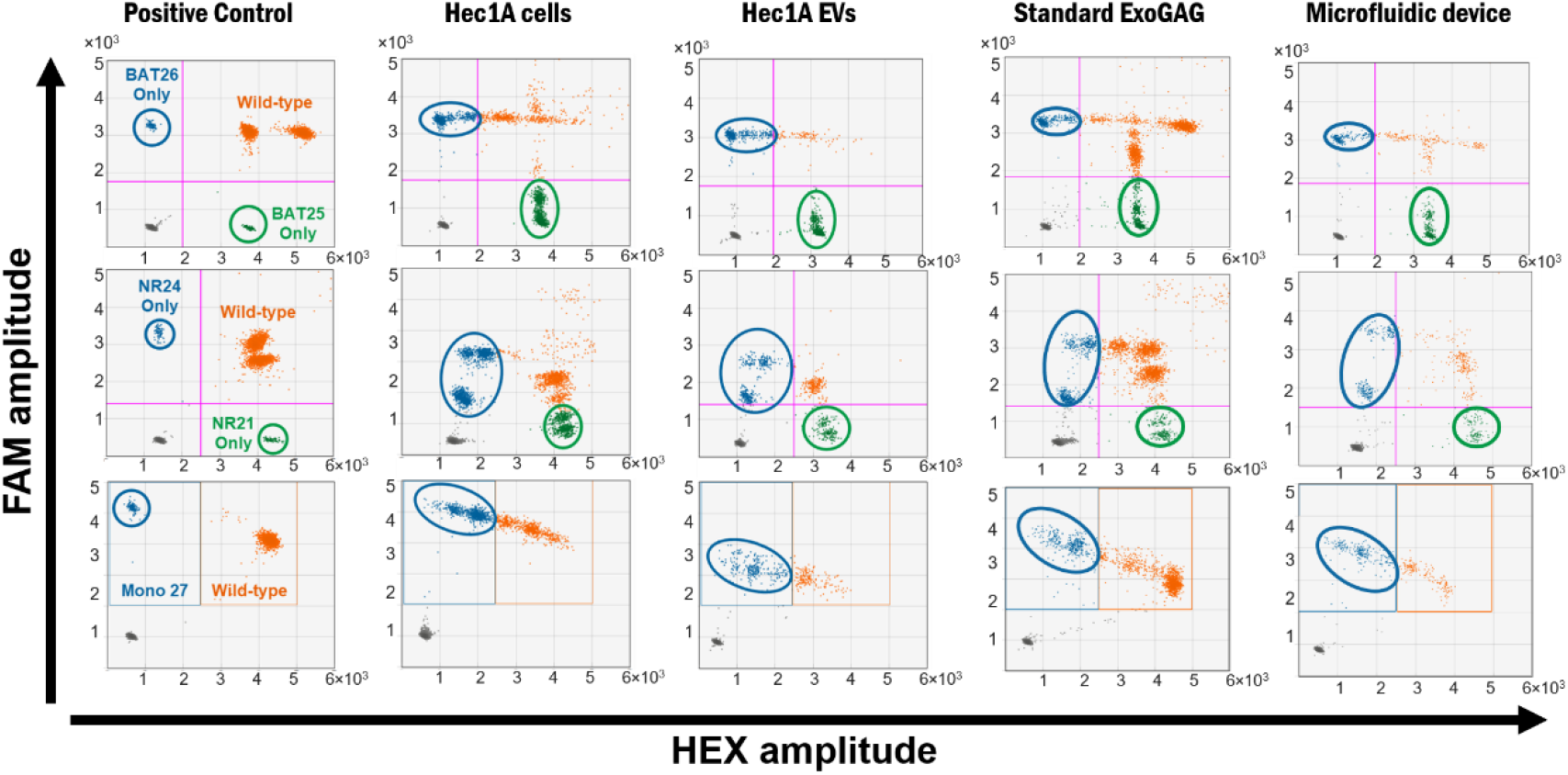
ddPCR analysis of MSI-associated markers in DNA recovered from Hec-1-A EV-spiked plasma. Five mononucleotide repeat markers were evaluated across three multiplex reactions: BAT25/BAT26, NR21/NR24 and MONO27

### Demonstration in clinical samples

#### MSI detection in endometrial cancer patients

Our long-term goal developing this method is to isolate EV DNA for detection of cancer mutations in liquid biopsies, which could lead to actionable clinical decision-making in patient care. While the method could be used for the detection of different nucleic acid markers, here we demonstrate it for the process of detecting MSI alteration in endometrial cancer (EC) patients. To ensure sufficient genomic material for downstream testing of cancer mutations in EV-DNA by EVNA-chip, we used between 0,5 to 2 ml of plasma as input volume for the pillar chip. We used ddPCR to detect MSI mutations. Our results demonstrate that the EVNA-chip workflow recovered sufficient EV-derived DNA from ExoGAG-enriched aggregates for downstream ddPCR analysis and enabled the detection of MSI alteration in plasma samples. The findings were aligned with those obtained using cell-free DNA, with MSI being detected in samples from MSI-high patients. Notably, clinical samples enriched from uterine aspirates showed superior performance, enabling MSI detection in cases where neither cell-free DNA nor EV-DNA analysis was able to identify MSI status.

#### miR-X detection in Fabry disease

To demonstrate our method ability to recover RNA cargo in a form suitable for downstream analysis of disease-associated targets we evaluated miR-X in samples from patients with Fabry disease. Fabry disease is a rare X-linked lysosomal storage disorder caused by deficient or impaired α-galactosidase A activity, leading to the progressive accumulation of glycosphingolipids and multisystem organ damage^[31]^. Our previous work identified altered circulating levels of miR-X in patients with Fabry disease and proposed this microRNA as a candidate biomarker requiring further validation. Since Fabry disease specific treatment provides greater benefit when initiated before irreversible organ damage occurs, biomarkers that may support earlier detection and disease monitoring are of clinical interest.

For microRNA analysis 2 ml of plasma was used as the input volume for the EVNA-chip workflow. The healthy donor controls presented miR-X concentrations of 7.33, 19.3, and 1.49 copies/µL, corresponding to a descriptive mean of 9.37 copies/µL. The Fabry disease samples presented concentrations of 1.97, 5.16, and 0.391 copies/µL, with a descriptive mean of 2.51 copies/µL (Figure 6). Although the limited sample size prevents statistical conclusions, these results demonstrate that the recovered RNA was suitable for reverse transcription and miR-X analysis by ddPCR.

**Figure 6.**
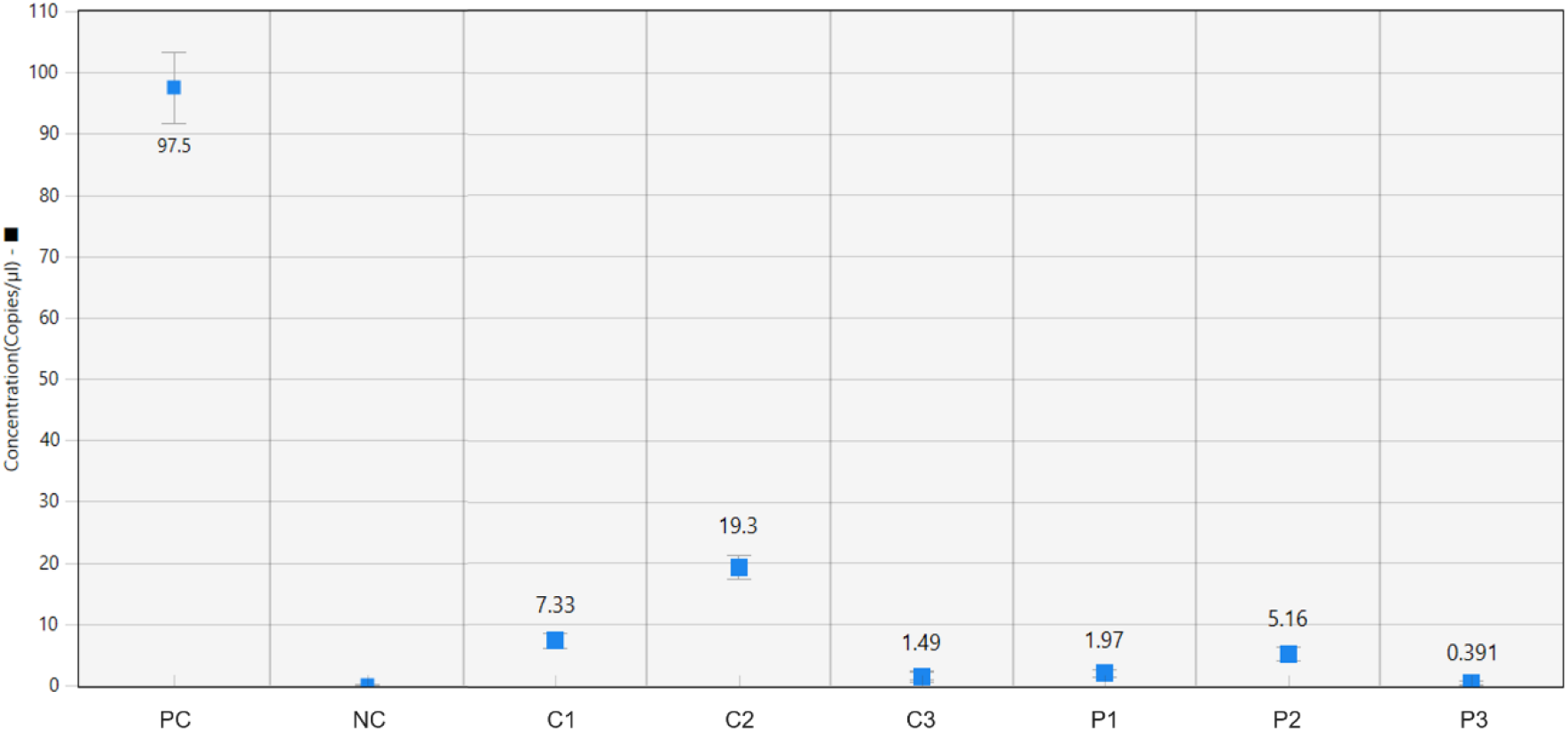
ddPCR quantification of miR-X in plasma samples from healthy donors and patients with Fabry disease. RNA recovered from plasma samples was reverse transcribed into cDNA and analyzed by ddPCR. Results are shown for the positive control (PC), negative control (NC), healthy-donor controls (C1–C3), and Fabry disease samples (P1–P3). Numerical labels indicate the measured miR-X concentrations in copies/µL, and error bars represent the confidence intervals calculated by the ddPCR analysis software.

## DISCUSSION

Here we demonstrate a method for EV nucleic acid extraction from biofluids, such as plasma and serum. This method combines ExoGAG-mediated aggregation of EVs and their mechanical entrapment in a microfluidic filtration chip (EVNA-chips), where they can be subsequently washed and lysed to collect the cargo. ExoGAG has been already established for EVs isolation but using standard centrifugation to precipitate the aggregates. Motivation to develop the microfluidic approach is to create a fully flow-based system, which would facilitate more compact automation using pumps and valves without robotics needed to automate centrifugation. Compared to the other microfluidic EV isolation methods, such as asymmetric flow field-fractionation (AF4) or nano-deterministic lateral displacement (nano-DLD) our device is simpler to fabricate and to operate, as each device is composed of only one microfluidic layer with minimum feature size of 10µm featuring one input and one output, where flow is driven by only one pump. Since the chip does not have any biomolecular functionalization, it does not need specific storage conditions. Similarly, use of ExoGAG instead of antibodies lowers the costs of reagents and facilitates their storage, though is also less specific. Microfluidic capture depends on the size distribution and amount of the aggregates. We found that aggregation would depend on the ExoGAG-sample ratio and on the type of biofluid, where plasma produced more abundant and larger aggregates compared to serum. To address these differences, dimensions of the microfluidic filter needed to be adjusted, which we achieved easily using modular approach, by stacking devices with three different designs in series. In order to evaluate the ability of the method to detect specific nucleic acid markers, we used a model where healthy serum or plasma samples were spiked with EVs featuring specific mutations, such as KRAS p.G12D and MSI from Hec-1-A cells. The yield of the nucleic acids isolation from both biofluids was about ∼40% (42.0 ± 4.28% of the EV-associated dsDNA from spiked serum and 40.7 ± 2.28% from spiked plasma. In the serum RNA experiments, recovery was 38.1 ± 1.46%) which was lower compared to the standard centrifugation-based approach, which had yield 94% or 47% for DNA or RNA, respectively. This difference may reflect losses at multiple stages of the workflow, including incomplete retention of smaller aggregates or adsorption of nucleic acids to the device material. Nevertheless, 0.55mL of spiked sample (about ∼ 5.14 x 10^9^ EVs from cells carrying mutation) was sufficient to be detectable in ddPCR, confirming that both yield, DNA integrity and possible interferences would not affect the downstream analysis. This is evidence that proposed method is well suited for assays to detect nucleic acid markers from EVs and could be basis for microfluidic automation that can reduce significantly time for manual handling.

The detection of KRAS p.G12D and MSI associated signal patterns demonstrated that DNA recovered through the microfluidic workflow remained amplifiable and compatible with mutation-specific and multiplex ddPCR assays. Mutant and wild-type droplet populations remained distinguishable after on-chip lysis and silica purification in Hec-1-A derived material and in spiked serum and plasma. These experiments provide evidence of downstream analytical compatibility.

The miR-X experiments extended the proof of concept beyond cancer-associated DNA. RNA recovered from patient-derived plasma could be reverse transcribed, and miR-X was quantitatively detected by ddPCR. The lower concentrations observed descriptively in the three Fabry disease samples were directionally consistent with the previous observation motivating this experiment. These findings demonstrate that RNA recovered using the workflow remains suitable for target-specific reverse transcription and ddPCR.

### Limitations of the study

Although ExoGAG provides an antibody-free enrichment strategy, further characterization of the recovered material will help clarify the potential co-enrichment of non-vesicular components and support optimization of sample purity. The present experiments do not distinguish interavesiclular NA from those associated with EV surfaces or co-isolated non-vesicular material. In addition, ExoGAG aggregation and silica-based nucleic acid purification are currently performed outside the device, providing opportunities for further integration through automated flow control and solution exchange. Finally, the successful downstream analysis of KRAS p.G12D, MSI associated alterations, and miR-X supports the analytical versatility of the workflow, while its diagnostic performance and clinical utility will require evaluation in future studies.

### Conclusions

Overall, this work establishes a modular platform for mechanically capturing EV aggregates and recovering nucleic acids through on-chip lysis. The recovery of amplifiable DNA and reverse-transcribable RNA supports the technical compatibility of the workflow with downstream ddPCR analysis. Future studies should focus on optimizing recovery, integrating automated flow control and solution exchange, expanding analytical characterization, and evaluating larger clinical cohorts to establish diagnostic performance and clinical utility. Once fully integrated and validated, the system could reduce hands-on time, operational complexity and costs associated with processing clinical plasma samples.

